# Microbial Worlds Apart: Distinct Communities in Crude Oil and Production Waters

**DOI:** 10.1101/2025.06.13.658281

**Authors:** Armando Alibrandi, Julia Plewka, Aurèle Vuillemin, Alexander Bartholomäus, Rolando di Primio, Alexander J. Probst, Jens Kallmeyer

## Abstract

Genomic analyses of microbial community composition are used to improve oil reservoir engineering and monitor reservoir dynamics. Given the challenges of extracting nucleic acids from oil, production water samples are often used as proxies from which to infer microbial community information from oil reservoirs. We employed 16S rRNA gene amplicon and metagenomic sequencing on samples of crude oil and production water from four North Sea oil fields. Taxonomic profiling revealed differences in microbial compositions and functions between production water and crude oil. Production water was more homogeneous, less diverse, harboured taxa associated with conditions non-native to the reservoir (e.g., seawater), and exhibited evidence of contact with atmospheric oxygen, most likely from passing through the water separators. Conversely, crude oil samples harboured microbial taxa typically associated with oil reservoirs. Despite long-term production and, in some cases, re-injection of production waters, the putative native microbial communities were still present in the oil. These findings demonstrate that crude oil samples are much more representative of oil reservoir microbiomes than their production water proxies.

## Introduction

Microbes have long been known to degrade hydrocarbons (Störmer, 1908), and such activity has been suspected to occur in subsurface oil reservoirs for many years (Bastin *et al*., 1926; Lipman and Greenberg, 1932; Zobell, 1945). In recent decades, these suspicions were confirmed through convergent observations, as highlighted by Magot *et al*. and references therein (Magot *et al*., 2005). In subsurface hydrocarbon reservoirs, microbial communities exert substantial influence on redox, mineralization dynamics, and other biogeochemical processes (Head, 2017). As long as temperatures remain below 80 °C (Wilhelms *et al*. 2001), most detrimental phenomena (e.g., oil souring, biodegradation) are driven exclusively by microbial activity (Connan, 1984; Huang and Larter, 2005; Magot *et al*., 2005; Medina-Bellver *et al*., 2005).

Prior to being able to mitigate specific microbial processes in oil or hydrocarbon reservoirs, one must first understand microbial community composition and the functional dynamics and transformative processes mediated by indigenous communities. Investigating microbial community structure in oil reservoirs requires the isolation of microbial cells from crude oil and/or production water, followed by nucleic acid extraction and sequencing analyses. Isolation of cells from production water can be achieved via simple filtration, and nucleic acids extracted directly from sample-laden filters. However, the isolation of cells from crude oil is much more challenging, invoking the use of non-polar solvents and/or non-ionic surfactants (Alibrandi *et al*., 2023; Dellagnezze *et al*., 2016; Sierra-Garcia *et al*., 2017; Yoshida *et al*., 2005).

During production, crude oil flows out of the reservoir to a production platform or Floating Production Storage and Offloading (FPSO) vessel. At this point, crude oil contains formation water naturally present in the reservoir and, if a breakthrough has occurred, injected water. In offshore reservoirs, typically seawater or separated formation water is injected into the reservoir to maintain pressure once its natural overpressure is depleted. As subsurface oil reservoirs are anaerobic, reducing environments, any microbial communities present are typically dominated by anaerobic organisms. In the production facility, however, separators remove water from the oil, and both of which interact with the ambient atmosphere and system infrastructure, e.g. pipes, storage tanks, *etc*. This exposure introduces microbial contaminants distinct from those resident in the reservoir and imposes a selective pressure favouring microbes capable of respiring oxygen. Microbes residing in oil reservoirs are thought to dwell in oil-water transition zones (OWTZ; Pannekens *et al*., 2019; Rajbongshi and Gogoi, 2021) or in water droplets suspended in the oil (Meckenstock *et al*., 2014). It is not unreasonable to assume that production water harbours microorganisms originating from the oil reservoir. However, direct contact of production water with oil-water separators, pipelines, injection fluids, and aquifers, is far more likely to contaminate production water samples than their purer oil reservoir counterparts.

Due to the convenience and simplicity of nucleic acid extraction from aqueous samples, examinations of microbial community structure in oil reservoirs have relied primarily on genomic analyses of production water samples. However, given their exposure to the atmosphere and processing infrastructure, we hypothesize that production water samples are less representative of a given reservoir’s oil-associated microbial communities than their crude oil counterparts. During water separation, production water is contaminated with microbes colonizing the separators and piping systems. In addition, production water mixes with other aqueous fluids (e.g., injection water, seawater, aquifers), introducing non-native taxa. In contrast, crude oil samples *sans* water separation are less prone to mixing with other fluids and are not exposed to oxygen in the separators, and as such, are more likely to retain the native microbial community composition.

To test this hypothesis, we analyse microbial communities in production water and crude oil samples and compare the representativeness of reservoir conditions. Employing 16S rRNA gene amplicon and metagenomic sequencing on samples from four North Sea oil fields, we report distinct differences in microbial community compositions and show that crude oil samples are more representative of the indigenous microbial populations in oil reservoirs. Our results bolster the current understanding of microbial dynamics and functions in oil reservoirs, clearly and convincingly demonstrating that crude oil and not production water should be used to monitor primary and secondary production, track fluid displacement, and identify non-native, transient microbes potentially deleterious to oil resources.

## Material and Methods

### Samples

Samples were provided by Aker BP (Lysaker, Norway), having originated from four North Sea oil fields located 200-300 km offshore of Stavanger (NO): Alvheim, Vilje, Volund, and Bøyla (Fig. 1 – Table 1). These consisted of both crude oil samples that did not undergo water separation and production water that was extracted from crude oil onboard the FPSO. For all four oil fields, *in-situ* reservoir temperature varied between 63°C and 66°C at producing depths ranging from 2,100 to 2,200 meters below seafloor (mbsf). Water depths ranged from 120 to 130 meters.

**Figure 1.**
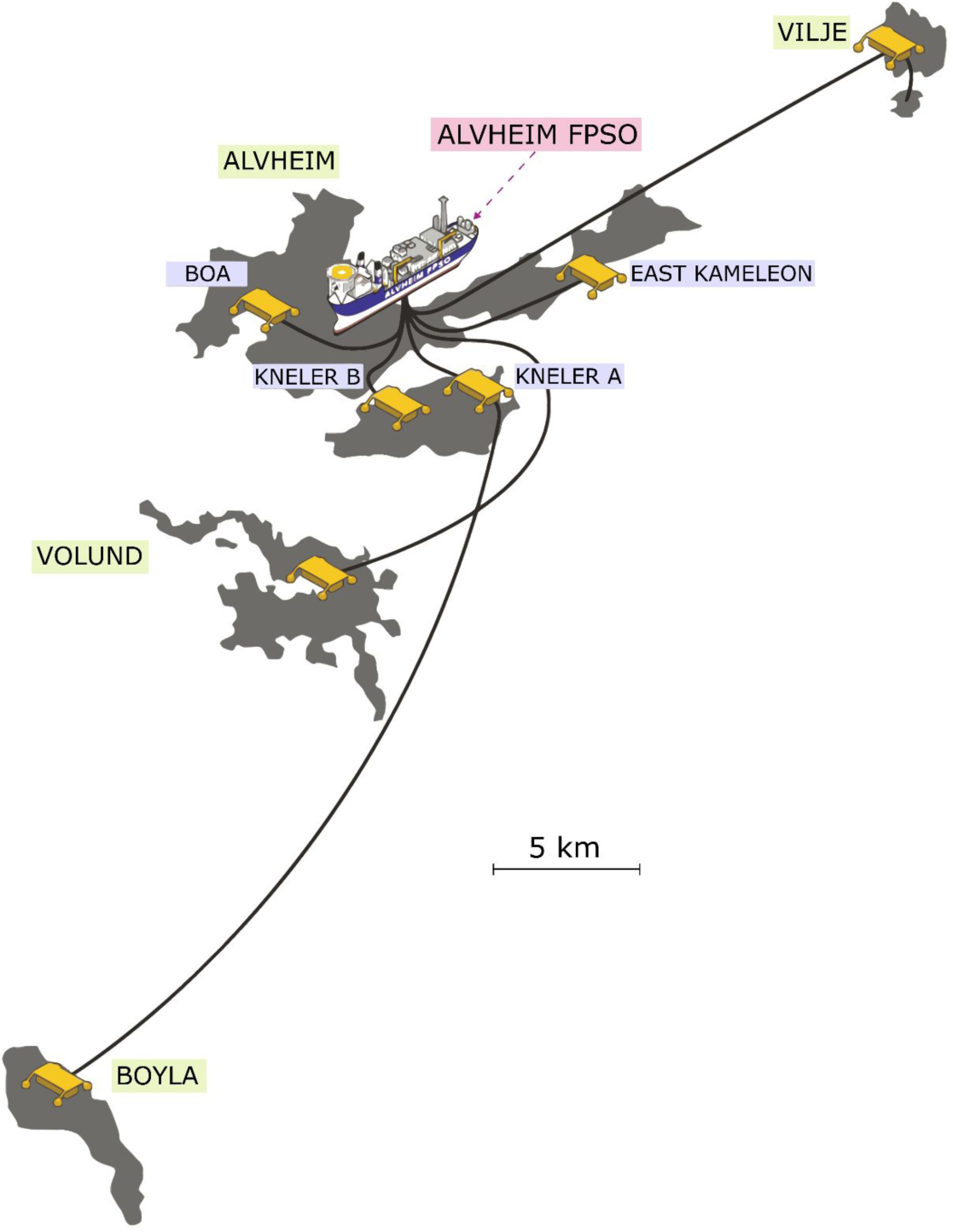
Map illustrating the distribution of four oil fields serviced by the Alvheim FPSO vessel (image adapted from AkerBP, 2024).

**Table 1.** Overview of the oil fields sampled with their respective year of discovery, commencement, and phase of production.

| *Field* | *Discovery*<br>*year* | *Production*<br>*start* | *Production method* | *Additional notes* |
| --- | --- | --- | --- | --- |
| *Alvheim* | 1998 | 2008 | Pressure maintained by underlying aquifer | Six discoveries including East Kameleon (E.Kam.), Boa, and Kneeler B (KNB) |
| *Vilje* | 2003 | 2008 | Pressure maintained by underlying aquifer | 20 km NE of Alvheim |
| *Volund* | 1994 | 2009 | Pressure maintained by underlying aquifer & water injection | 10 km south of Alvheim |
| *Bøyla* | 2009 | 2015 | Pressure maintained by water injection | 28 km south of Alvheim |

Alvheim oil field comprises six discoveries, three of which were sampled for this study: East Kameleon (E. Kam.), Kneeler B (KNB), and Boa. The oil field was discovered in 1998 and production commenced in 2008. The field is serviced by subsea production wells connected to the Alvheim FPSO vessel, and production is maintained by water pressure exerted by an underlying aquifer.

Vilje, Volund, and Bøyla represent distinct oil fields tied to the Alvheim FPSO. Vilje was discovered 20 kilometres northeast of Alvheim in 2003 and has been in production since 2008. This field features three horizontal subsea wells linked to the Alvheim FPSO. Oil is produced via water pressure exerted by the underlying aquifer. Volund was discovered 10 km south of Alvheim in 1994 and production started in 2009. This field exploits significant pressure from the aquifer sustained by injection of produced water delivered from the Alvheim FPSO. Bøyla was discovered 28 km south of Alvheim in 2009 and production began in 2015. Production in this field requires water injection to sustain pressure in the reservoir, supplemented with gas lift for well flow support.

Samples were collected from each of the Volund, Bøyla, Vilje, E. Kam., Boa, and KNB reservoirs and oil fields (Fig. 1). All samples were collected aboard the Alvheim FPSO vessel. Crude oil samples were collected directly from wellheads prior to water separation, and production water samples were obtained from separators following phase separation. All samples were transferred to sterile, PTFE-capped glass bottles without headspace to maintain anaerobic conditions, kept at room temperature during transport, and stored at 4 °C at GFZ in Potsdam until analysis.

Upon arrival at GFZ, samples were opened and subdivided into reaction tubes in an anaerobic glove box to avoid oxygen contamination. Samples were stored with a nitrogen headspace to ensure strict anoxia and avoid the growth of microaerophilic organisms. Although crude oil is composed of oil and a minor fraction of water, for the sake of simplicity we will henceforth refer to these samples as “oil.” All samples were delivered in duplicate bottles, each of which was subdivided into duplicate aliquots, resulting in four biological replicates per sample.

### DNA extraction

In an anaerobic glove box, oil samples were mixed 1:1 with sterile phosphate-buffered saline (PBS) containing 0.1 vol % Tween 80. Following vigorous shaking, samples were stored overnight at 55 °C. The next day, the polar fraction of the solution was filtered through Sterivex filters (pore size 0.22 µm; Merck KGaA, Darmstadt, Germany). To avoid contamination with airborne microbes, handling of the filter and subsequent DNA extraction was carried out in a laminar flow cabinet. Filter casings were cracked open, and filters were transferred to bead tubes. Prior to continuing with the manufacturer’s protocol for the PowerSoil Pro DNA extraction kit (DNEasy Power soil pro kit, Qiagen, Hilden, Germany; Cat. No. 47014), 40 µl of 10 % sodium dodecyl sulphate (SDS) was added to the bead tubes. The addition of SDS increased DNA yield from crude oil samples (Alibrandi *et al*., 2023). We opted for this extraction protocol over the isooctane method (Alibrandi *et al*., 2023) to ensure comparability between oil and production water samples. Despite yielding less nucleic acid than the isooctane method, the Tween 80-PBS technique provided cleaner DNA and facilitated subsequent handling with the 2 ml bead tubes of the extraction kit of the Power Soil Pro Kit. With respect to microbial community structure, parallel tests were performed to ensure reproducibility and consistency between the isooctane and surfactant-based DNA extraction methods.

Production water samples were incubated overnight at 55°C (consistent with the oil samples), filtered, and processed as described above. DNA extractions on blank Sterivex filters were conducted as negative controls. DNA was quantified using a Qubit 2.0 device and a dsDNA HS assay cocktail containing buffer and a fluorescent dye (Invitrogen, Carlsbad, USA).

### Amplification 16S rRNA genes and amplicon sequencing

All samples were processed in triplicate. Bacterial and archaeal 16S rRNA gene fragments (V4 hypervariable region) were PCR amplified with the universal barcoded primer pair 515F (5′-GTG TGY CAG CMG CCG CGG TAA-3′) and 806R (5′-CCG GAC TAC NVG GGT WTC TAA T-3′). The final volume of each reaction mixture was 50 µl, consisting of 2 µl DNA template, 0.5 µl Taq DNA polymerase, 2 µl dNTP, 2 µl MgCl_2_, 5 µl 10 × polymerase buffer, 0.5 µl BSA, 2.5 µl of each primer, and 33 µl PCR grade water (EurX, Gdansk, Poland). PCR amplification was run at 95 °C for 5 min of initial denaturation, followed by 32 cycles of 30 sec at 95 °C (melting), 30 sec at 56 °C (annealing), and 1 min at 72 °C (elongation), with a final elongation of 7 min at 72 °C. PCR products were cleaned using AMPure magnetic beads (Beckman Coulter, Brea, USA), and barcoded samples were normalized to 20 ng of DNA and pooled. Amplicon sequencing was performed on an Illumina MiSeq platform using 2 × 300 base pair (bp) reads at Eurofins Genomics (Ebersberg, Germany).

### 16S rRNA gene sequencing data treatment and statistical analyses

Read demultiplexing was performed using Cutadapt v. 3.5 (Martin, 2011) with the following parameters: –e 0.2 –q 15,15 –m 150 –-discard-untrimmed. Amplicon sequence variants (ASVs) were generated using trimmed reads and the DADA2 package v. 1.20 (Callahan *et al*., 2016) with R v. 4.1, applying the pooled approach with the following parameters: truncLen = c(220,180), maxN = 0, rm.phix = TRUE, minLen = 160. Taxonomic assignment was performed using DADA2 against the SILVA 16S rRNA SSU database release 138 (Quast *et al*., 2012). ASVs representing chloroplasts, mitochondria, and/or singletons were removed. Partial 16S rRNA gene sequences were aligned using SINA online v.1.2.11 (Pruesse *et al*., 2007).

Statistical analyses of alpha and beta diversity were conducted using the Vegan community ecology package in RStudio v. 4.3.1 and PAST 4.14 software (Hammer *et al*., 2001). The 16S rRNA gene sequence dataset was rarefied to 1,500 reads, and all downstream statistical analyses were performed using this rarefied dataset. Two metrics were applied to evaluate microbial composition: relative read abundance of ASVs and α-diversity, including observed richness and Shannon indices. To estimate regional differences across oil fields and reservoirs, we compared the beta-diversity of samples based on their microbial compositions, conducted Principal Coordinate Analyses (PCoA) with the Bray-Curtis similarity index, and performed non-parametric Permutational Multivariate Analyses of Variance (PERMANOVA). The results of PCoA presented dissimilarity between microbial communities, providing insights into clustering and other trends across regions, while the results of PERMANOVA were used to determine the statistical significance of observed differences.

Indicator Value (IndVal) analyses (Dufrêne and Legendre, 1997) were conducted to identify taxa indicative of either production water or oil samples. This method considers both relative species abundance and occurrence frequency to determine the specificity and fidelity of a given group. IndVal scores and associated p-values were calculated to assess the significance of taxa as indicators, with a threshold p-value of 0.05 applied to determine statistical significance (supplementary table S8). We then divided all available sample replicates into two groups, production water (n = 22) and oil (n = 28), and performed IndVal analyses using 16S rRNA gene sequence data (Supplementary Fig. S1).

### Metagenomic sequencing, de novo assembly, and gene annotation

DNA extracts were sent to CeGaT GmbH (Tübingen, Germany) for metagenomic sequencing. Libraries were prepared using the Nextera XT DNA Library Preparation kit (Illumina), and sequencing was performed on a NovaSeq 6000 Illumina platform at CeGaT, aiming for 50 million read pairs (2 × 150 bps).

### Quality control and assembly of shotgun metagenomic reads

BBDuk (v. 37.09; Bushnell, 2014) was used to remove Illumina artefacts and adapters from raw reads, quality control and trimming was accomplished using Sickle v. 1.33 (Joshi and Fass, 2011), and reads were de-duplicated with BBMap (v. 37.09; Bushnell, 2014). Sourmash (v. 4.8.7; Pierce *et al*., 2019) was run on quality-checked forward reads with a *k*-mer (short sequence of length *k*) of size 31 to calculate Jaccard similarities between metagenomic samples. A Jaccard similarity of one indicates identical samples, while zero means that no identical *k-mers* are found in the read sets. Quality-checked reads were assembled with metaSPAdes (v. 3.15.5; Nurk *et al*., 2017), and we then removed all scaffolds smaller than 1,000 bp from FASTA files with pullseq (v. 1.0.2; Thomas *et al*., 2015; github.com/bcthomas/pullseq). Prodigal (v. 2.6.3; Hyatt *et al*., 2010) was run in meta mode to predict and translate genes into amino acid sequences.

Functional annotations of predicted open reading frames (ORFs) were performed using the DIAMOND protein aligner v. 0.9.24 (Buchfink *et al*., 2015), and proteins were annotated with eggNOG v. 2.1.12 and eggNOG DB v. 5.0.2; Cantalapiedra *et al*., 2021; Huerta-Cepas *et al*., 2019). We then performed a quantitative functional annotation, focusing on ORFs encoding proteins associated with sulphate, sulphite, and polysulphide reduction, nitrate and nitrite reduction, hydrocarbon degradation, methanogenesis, heat shock response, salt stress, biofilm formation, and/or microaerophilic respiration (Supplementary Tables S2-S3). A Wilcoxon paired test was run to assess whether metabolic-associated ORFs differed significantly between oil and production water samples. Finally, we calculated delta values and ran one-sample t-tests to determine whether the differences between these sample types were significant.

### PCoA of functional gene sequences

The 228,972 nucleotide sequences belonging to predicted genes from all metagenomes were clustered with MMSeqs2 (v. 15.6f452; Steinegger and Söding, 2017) in cluster-mode 2, coverage-mode 1, with a minimum breadth of 95% and minimum sequence identity of 95%. Quality-checked reads were mapped against representative sequences from the resulting 77,195 clusters with bowtie2 (v. 2.3.5.1; Langmead and Salzberg, 2012) in sensitive mode. Mappings were filtered to exclude all reads with more than five mismatches, and sequences were deemed present in samples if their minimum coverage breadth was greater than 95%.

Coverage values were normalized based on the bp counts of the forward read sequences. Only gene sequence clusters present in more than one sample were used for the PCoA calculation (to preclude the influence of under-sampling; 36,831 non-singleton clusters remained). Bray-Curtis distances were calculated from the normalized mean coverage depth using the R vegan package (Anon, 2024). MRPP and Adonis tests were run in R (R Core Team, 2020; RStudio Team, 2020) to identify significant differences between sample types.

### Prokaryotic community composition based on extended rpS3 gene sequences

The ribosomal protein S3 marker gene (*rpS3*) was used to estimate the extent of diversity in prokaryotic community composition in metagenomes. Compared to 16S rRNA gene analyses, *rpS3*-gene-based diversity analyses provide higher phylogenetic resolution, greater variability, and reduced susceptibility to horizontal gene transfer, making it a more reliable tool for resolving taxonomic assignments (Sharon *et al*., 2015). Marker genes were identified via species-specific Hidden Markov Models (HMMs) and by comparing the amino acid sequences of predicted genes with diamond BLASTp (e-value cut-off of 1e-5; Buchfink *et al*., 2015) against the UniRef100 database (downloaded on 23.06.2021; The UniProt Consortium, 2019). The *rpS3* gene nucleotide sequences with 1,000 bp flanking regions were extracted from all samples and clustered with MMSeqs2 (v.15.6f452; Steinegger and Söding, 2017) in cluster-mode 2, coverage-mode 1, with a minimum breadth of 95 % and minimum sequence identity of 95 %.

*RpS3* sequences were taxonomically annotated by comparing them with *rpS3* gene sequences extracted from the GTDB database (GTDB v. 214; Parks *et al*. 2022) with usearch – ublast (v. 10.0.240_i86linux64; Edgar, 2010), detailing corresponding coverage, breadth, and taxonomy of extended *rpS3* gene sequences (Supplementary Table S5). Quality-checked reads from all samples were mapped against representatives using bowtie2 (v. 2.3.5.1; Langmead and Salzberg, 2012) in sensitive mode. Reads with more than five mismatches were excluded. Mean coverage depths of extended *rpS3* gene sequences were calculated for all sequences with a coverage breadth greater than 95%. ANOVA and Tukey HSD tests were run in R (R Core Team, 2020; RStudio Team, 2020) to identify significant differences in community composition between oil and production water samples.

### Binning of metagenomic assembled genomes (MAGs) and genome comparison

Length-filtered scaffolds were binned into MAGs using Abawaca v. 1.0.0 (Brown *et al*., 2015) and MaxBin2 v. 2.2.7 (Wu *et al*., 2016) with default parameters, bins were aggregated with DASTool v. 1.1.6 (Sieber *et al*., 2018), and the resulting selection was manually curated with uBin v. 0.9.14 (Bornemann *et al*., 2023). We then calculated completeness and contamination with CheckM2 v. 1.0.1 (Chklovski *et al*., 2023) and used GTDB-tk v. 2.4.0 (Chaumeil *et al*., 2019) with the Genome Taxonomy Database v. 214 (Parks *et al*., 2022) to assign taxonomy. The resulting taxonomic annotation was used to construct a *de novo* tree of MAGs with GTDB-tk v. 2.4.0 (Chaumeil *et al*., 2019) de_novo_wf workflow, converted into an iTOLs usable format with GTDB-tk’s convert_to_itol workflow, and visualised with iTOLs (v. 6; Letunic and Bork, 2024). We then applied Bowtie2 v. 2.4.1 (Langmead and Salzberg, 2012) and samtools v. 1.13 (Li *et al*., 2009) to map all quality-checked reads against the combined MAG from all samples. Mappings were used to predict strain diversity and compare MAG presence/absence between oil and production water samples via inStrain v. 1.8.1 (Olm *et al*., 2021).

Coverage values of strains extracted from MAGs were normalised by the bp count of the quality-controlled reads, to adjust for different sequencing depths for oil vs. water samples. Normalised coverages, though estimates of abundances, were not converted to relative abundances. The low abundance human contaminant *Streptococcus oralis_S,* (Ren *et al*., 2024) was discarded for further analysis (inStrain coverage and breadth values in samples provided in Supplementary Table S7). Figures displaying resulting data were constructed in R v. 4.2.2 (R Core Team, 2020) and RStudio v. 2023.09.0+46; RStudio Team, 2020) using the ggplot2 package (Hadley Wickham, 2016).

## Results and Discussion

### Microbial communities in oil and production water

Ribosomal RNA gene analyses identified ASVs typically associated with hot oil reservoirs (*i.e.* close to 80°C, Wilhelms *et al*., 2001), including species of *Caminicella*, *Petrotoga*, and the halophilic *Flexistipes* genus (Fig. 2A; Christman *et al*., 2020; Dahle *et al*., 2008; Vigneron *et al*., 2017). Data from 11 metagenomic libraries (five oil and six production water samples) supported these findings, revealing distinct taxonomic and genetic diversity between oil and production water samples (Fig. 2C, supplementary table S5).

**Figure 2.**
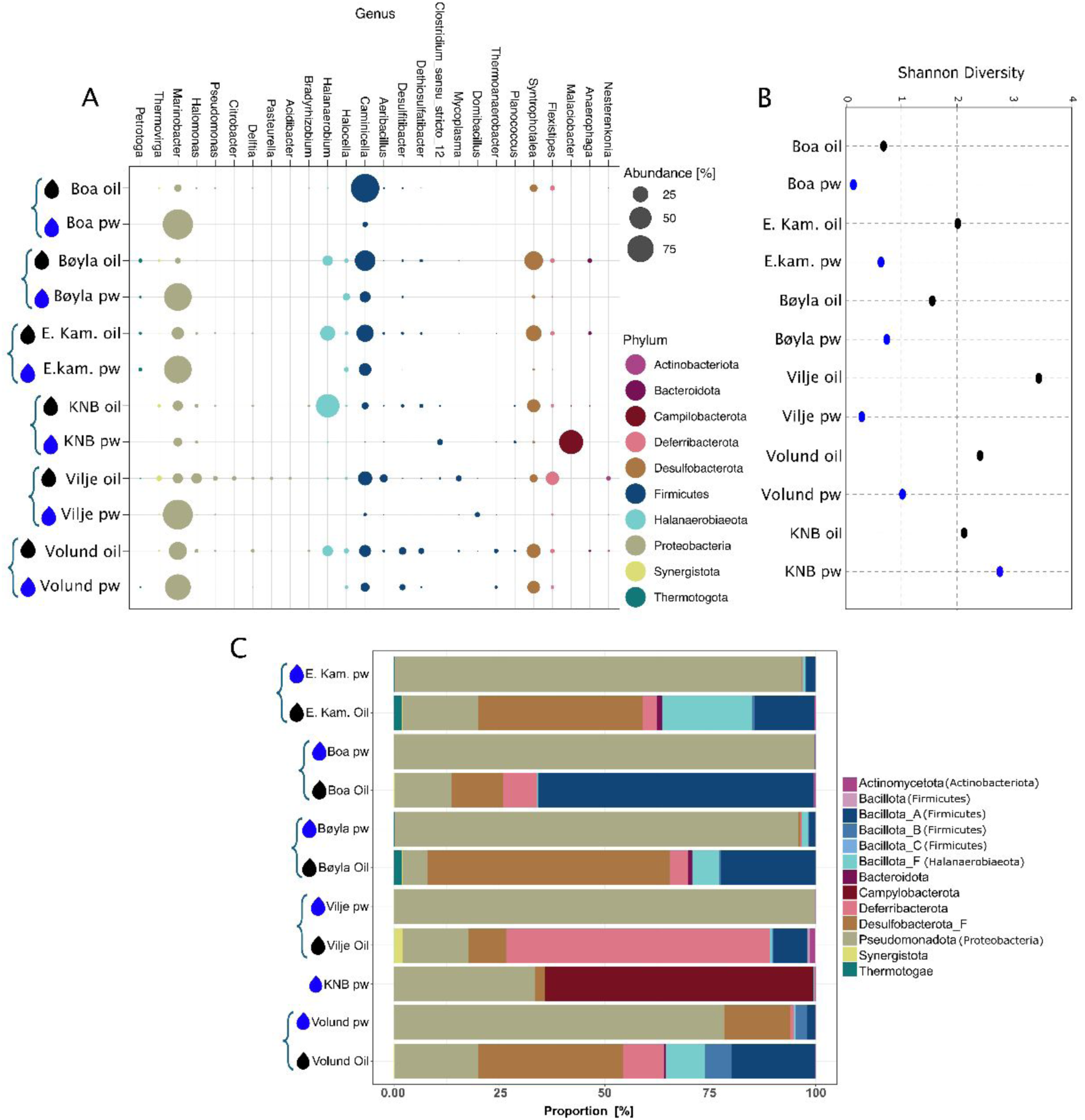
Taxonomic affiliation and diversity found in oil and production water samples. (A) Bubble plot of the relative abundances of the 98% most abundant taxa (n = 4), taxonomically annotated with DADA2 against the SILVA 16S rRNA SSU database (release 138; Quast *et al*., 2012). (B) Shannon diversity indices of oil and production water (pw) samples (n = 4). (C) Community composition of prokaryotic organisms in the eleven samples based on coverage of extended *rpS3* gene sequences. Different colours denote various proportions of different phyla. Taxonomy was assigned by comparing against a database of *rpS3* sequences extracted from the GTDB, along with the respective taxonomies (GTDB v. 214; Parks *et al*., 2022).

Taxonomic profiling based on the *rpS3* marker gene identified oil reservoir-associated taxa, including *Haloanaerobium congolense* (Ravot *et al*., 1997), *Petromonas tenebris* (Christman *et al*., 2020), *Petrotoga* spp. (Miranda-Tello *et al*., 2004), *Thermoanaerobacter thermocopriae*, and members of the *Thermovirgaceae* family (Dahle and Birkeland, 2006). Production water samples were dominated by seawater-inhabiting aerobic genera, such as *Marinobacter* spp. (Xu *et al*., 2008), which accounted for up to 98% of the 16S rRNA-inferred relative abundance in the Boa reservoir, and *Malaciobacter marinus* (Miller *et al*., 2018), which accounted for 61% of the relative abundance in KNB production water. Taxa native to the reservoir were either absent or underrepresented in production water samples. For instance, species of *Caminicella* accounted for 86 and 2% of the relative abundance in crude oil and production water in the Boa reservoir, respectively. *Syntrophotalea* spp. accounted for 6% of the relative abundance in the Vilje reservoir’s oil but were undetected in its production water. This trend was consistent across all reservoirs.

Oil samples exhibited greater microbial diversity than production water samples, harbouring a mean of 2.8 times more phyla (Supplementary Table S5). Bacillota_A (p-value 0.009) and Desulfobacterota_F (p 0.0042) were significantly more abundant in oil samples, while Pseudomonadota (p 1.1*10^-11^) were detected in greater abundance in production water. Shannon diversity indices derived from 16S rRNA gene sequencing data (Fig. 2B) supported these findings, indicating greater diversity in oil samples. These results oppose previous findings by Cai *et al*. (2015), who reported more distinct OTUs in production water than in oil. The Vilje reservoir yielded the most expansive microbial diversity, while Boa displayed the most conserved. The observed differences in diversity between production water and oil likely reflect the inability of seawater-associated taxa to survive harsh reservoir conditions.

PERMANOVA analyses (Supplementary Table S1) showed significant differences across most sample groups, with a few exceptions. E. Kam. oil did not differ significantly (p > 0.05) from that of Bøyla, Vilje, or Volund, perhaps due to the central location of E. Kam. In a similar vein, Volund production water was comparable to its corresponding oil, and the production waters from Vilje and Boa showed no significant differences. These reservoirs maintain pressure via underlying aquifers, suggesting that the production waters’ microbial composition may be influenced by aquifer-derived taxa. As a polar fluid, formation water mixes much easier with aquifer water than non-polar oil. This suggests that production water is more susceptible to environmental influences than oil, potentially limiting its ability to reflect the native reservoir microbiome. However, distinct microbial signatures observed in most reservoirs highlight the potential of microbial community variation as a valuable proxy for tracing oil migration and subsurface distribution (Zhang *et al*., 2020, 2021).

We reconstructed 26 MAGs from oil samples and 21 from production water samples (Supplementary Fig. S2). Production water MAGs were of higher quality (16 of 21 > 95% completeness, < 5% contamination) than their oil counterparts (8 of 26 were high-quality; Supplementary Fig. S3). This is most likely due to hydrocarbon interference in DNA extraction from oil samples. The greater community complexity in oil samples, as indicated by higher Shannon indices (Fig. 2B), may also contribute to the lower quality of oil MAGs. The greater number of sequenced base pairs in production water samples (mean bp count 1.6 times higher than in oil samples; Supplementary Table S4), could also play a role in this disparity.

Strains of anaerobic, halophilic, and thermophilic taxa, such as *Petromonas tenebris*, members of order *Halanaerobiales*, and *Flexistipes* spp. (Fig. 3; Supplementary Fig. S2) were present both in production water and oil but were substantially more abundant in the latter (mean difference = 10.6, 9.9, and 18.3-fold, respectively). Species of *Marinobacter* were abundant in all production water samples, while they comprised but a minor fraction of communities in oil samples (mean 6.4-fold lower than production water; not detected at all in the Bøyla sample). Hence, these strains likely originated from water that was injected into the reservoir and, as such, are more likely contaminants (either as cells or nucleic acid remnants) than authigenic elements of the *in-situ* oil communities.

**Figure 3.**
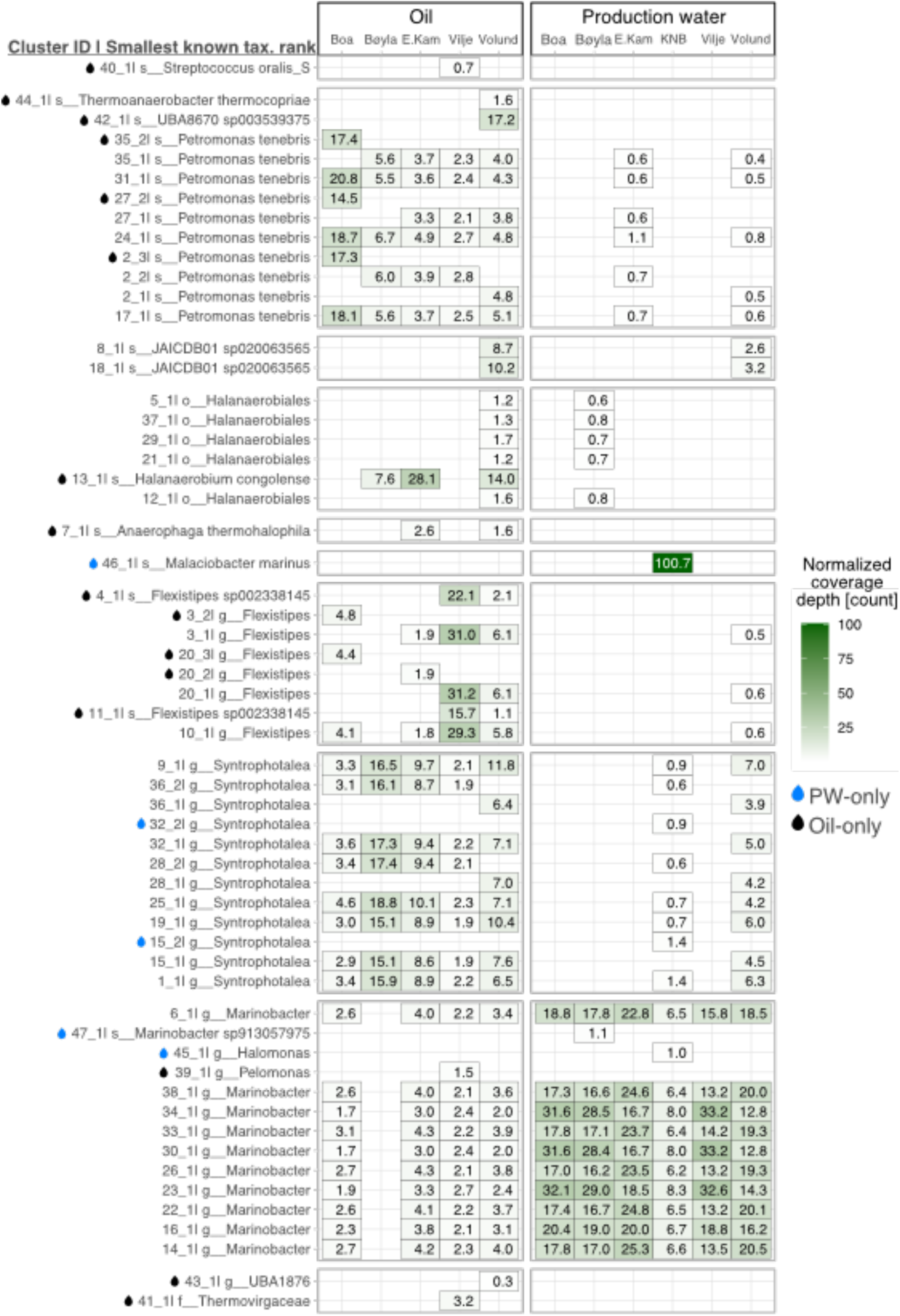
Normalised coverage depth of strains extracted from eleven oil and production water samples. Normalised coverage depth [count] of strains identified in MAGs assembled from five oil and six production water metagenomes. Normalised coverage was calculated by adjusting the mean coverage depth of the entire MAG for differences in sequencing depth (base pair counts of the raw reads) of the metagenomes. This facilitated a direct comparison between oil and water samples. Clustering IDs consist of the species and strain number from InStrain (Olm *et al*., 2021). Genomes are grouped by phyla and named after the smallest known taxonomic rank annotated through GTDB-tk (Chaumeil *et al*., 2019). Strains present exclusively in oil or water samples are marked in black and blue, respectively.

While most strains were present in at least one production water sample and oil sample of varying normalised coverage depth, five strains belonging to the genera *Syntrophotalea* (2), *Malaciobacter* (1), *Marinobacter* (1), and *Halomonas* (1) were only present in production water samples. As such, we conclude that these organisms do not originate in the reservoir but are contaminants from the production process. By comparison, 16 strains, the majority belonging to phyla *Bacillota_A*, *Deferribacterota,* and *Synergistota*, were present exclusively in oil samples. These strains likely originated from the reservoir and appear refractory to transfer into aqueous environments (Cai *et al*., 2015).

We determined indicator species for both production water and oil samples based on *16S rRNA* gene sequence data. The abundance of several *Marinobacter* ASVs yielded high IndVal scores (> 80%) and correlated significantly (p < 0.0001) with production water samples. In contrast, species of *Caminicella*, *Synthrophotalea*, *Haloanaerobium*, *Halomonas*, *Flexistipes,* and *Thermovirga* yielded IndVal scores above 75% and correlated significantly (0.0001 < p < 0.0003) with oil samples (Supplementary table S8). While oil and production water harboured different microbiomes, the microbial communities in production water appear to imprint on the oil communities, and vice versa, as evidenced by the co-occurrence of several strains across both sample types. Genomes constructed from oil and production water samples exhibited marked variations in normalised coverage, and significant differences in indicator species profiles confirm substantial differences between these ecosystems (Fig. 2).

### Functional adaptation to life in oil reservoirs

We evaluated protein groups relevant to life in oil reservoirs and found major differences in the metabolic potential between habitats (supplementary figure S4). Methanogenesis-related ORFs were rare and detected only in oil samples from Volund (5), Vilje (1), and E. Kam. (1). Volund yielded the fewest ORFs related to microaerophilic respiration. Microaerophilic respiration was evident across oil and water samples, and particularly abundant in production water samples from Vilje and Boa. While evidence posits both oxygen consumption and production in deep seafloor environments and/or deep subsurface groundwater (Ruff *et al*., 2023; Sweetman *et al*., 2024), the presence of oxygen respiration-related ORFs does not imply that oxygen is present in the oil reservoirs – rather that taxa bearing these ORFs might be able to cope with oxygen stress.

Osmotic stress-related ORFs were more abundant in oil than in production water, which supports our hypothesis that oil samples more accurately reflect the true microbial community composition of the reservoir. Sulphate and nitrate are microbially reduced in oil reservoirs and thus depleted rather quickly (Vigneron *et al*., 2017). The presence of ORFs specific to dissimilatory sulphate and nitrate reduction in all samples suggests that the oil fields experience replenishment of the electron acceptors sulphate and nitrate via injection of sulphate-rich seawater (2.7 g kg-1) and/or from underlying aquifers. Upon conducting a Wilcoxon paired test (supplementary table S3), we did not observe statistically significant differences (p < 0.05) between single metabolic processes in oil and production water. However, a marginally significant trend (p = 0.059) was observed for nitrate reduction, osmotic stress, and microaerophilic respiration ORFs. To further evaluate differences across metabolic pathways, we conducted a one-sample t-test on delta values between oil and production water samples. A significant effect (p 0.001) indicated a shift in metabolic activity favouring oil-associated samples. This pattern is consistent with the findings of Cai *et al*. (Cai *et al*., 2015) and suggests that while individual pathways may not show strong statistical separation, there is a metabolic distinction between the two environments.

### Genetic distance between production water and oil reservoir communities

Distinct clustering of microbial communities originating from oil or production water samples was confirmed by Bray-Curtis distances based on normalized 16S rRNA gene coverage values, coverage of gene sequence clusters recovered from the eleven metagenomes (Fig. 4A-B), and Jaccard similarities of the complete quality-checked read set (Fig. 4C). While Jaccard’s similarity might be influenced by sequencing depth, it has been previously used to differentiate metagenomic samples from different sites (Bozzi *et al*., 2024). The highest intra-metagenomic Jaccard similarities resulted between Boa and Vilje production water samples, ranging from 0.14 to 0.197 (Fig. 4C). This might be due to the fact that both of these reservoirs produce oil solely via pressure exerted by underlying aquifers, *sans* extraneous. PCoA plots also display production water samples in clustered arrangements (Fig. 4A-B), indicating a high degree of similarity across all samples.

**Figure 4:**
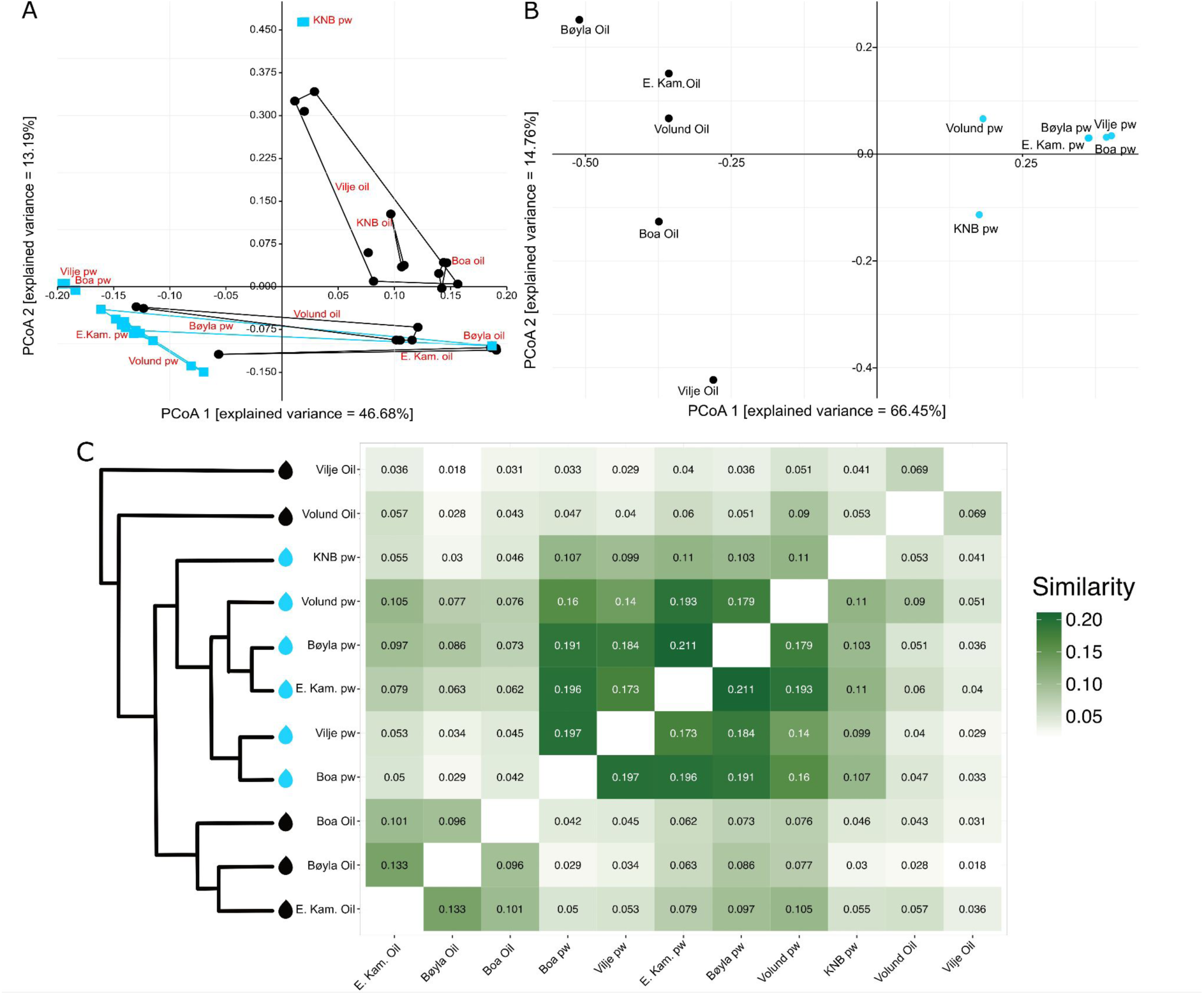
PCoA and Jaccard distances within oil and production water samples. (A) PCoA plot based on normalised coverage of 16S rRNA gene sequences showing dissimilarity between samples. Oil samples are black circles and production water samples are blue squares (n = 4). (B) PCoA plot based on the abundance of 36,831 gene sequence clusters in eleven samples (oil samples in black, production water samples in blue). (C) Heat map and dendrogram of Jaccard similarity and clustering between quality-checked metagenomic reads calculated by sourmash.

Oil samples are dispersed more widely, indicative of substantial variability and consistent with observations that oil samples are more diverse than water samples (Fig. 2B). In addition to the aforementioned reasoning regarding harsh conditions in oil reservoirs necessitating different survival strategies, pristine communities in geologically distinct reservoirs are also known to differ due to isolation and varying abiotic factors (e.g., temperature, pH, salinity; Gao *et al*., 2016). The distance between gene sequence clusters of oil samples from the Vilje and Bøyla reservoirs mirrors their geographical distance (Fig. 4B), underscoring the potential utility of genomic analyses in oil field monitoring (Zhang *et al*., 2020).

Jaccard similarity scores between quality-checked reads from oil and water samples collected from the same site are relatively low, ranging from 0.09 (Volund) to 0.029 (Vilje; Fig. 4C). This is indicative of samples that are not highly similar. Since we could identify the same strains in reservoirs having different lithologies (Fig.3), we compared water and oil samples from different sites. Resulting high similarity scores between unrelated oil and water sample pairings (e.g., Volund prod. Water vs. E. Kam. oil; Jaccard similarity = 0.105), suggest that production water and oil do share a measurable suite of identical *k-mers*. We conclude that the similarity between production water samples is greater than the similarity between oil and production water.

Multi-Response Permutation Procedures (MRPP; significance of delta = 0.005) and Adonis tests (R² value of community type = 0.62; p 0.03) based on the coverage of gene sequence clusters recovered from the eleven metagenomes confirmed that observed differences are indeed significant. The chance-corrected within-group agreement (A of MRPP) was 0.3753 (based on observed delta = 0.3678 and expected delta = 0.5888), which suggests that observations within the two groups are much more alike than would be expected by random grouping. This confirms that the microbial communities of oil and production water samples indeed represent different ecosystems. These findings reinforce the notion that oil and production water sustain fundamentally different microbial communities, despite some degree of cross-influence.

### Conclusion

Distinct differences in taxonomy, genetic diversity, and adaptation patterns between oil– and production water-associated microbial communities suggest that the latter are heavily contaminated with non-indigenous microorganisms. As such, we conclude that production water can serve neither as a proxy for microbial community composition nor functional capacity of the oil reservoir environment. Although the Alvheim oil field has been in production for many years, with the most recent production commencing ten years ago (2015), the taxa present in its oil samples suggest that at least a fraction of its native microbiome has not yet been overprinted by transient contamination. Taken collectively, the findings of this investigation necessitate a paradigm shift in sampling procedures for accurate profiling and functional analyses of the microbial communities in oil reservoirs, prioritising crude oil samples that have not undergone separation processes over production waters.

## Notes

The authors declare no conflict of interest.

## ACKNOWLEDGMENT

A special thanks go to my colleagues and technicians at GFZ who helped materialise this work and to my family for the immense support.

