## Supplementary figure for "Microbial Worlds Apart: Distinct Communities in Crude Oil and Production Waters"

(3) Aker BP, Norway.

**Content:** supplementary figures and tables

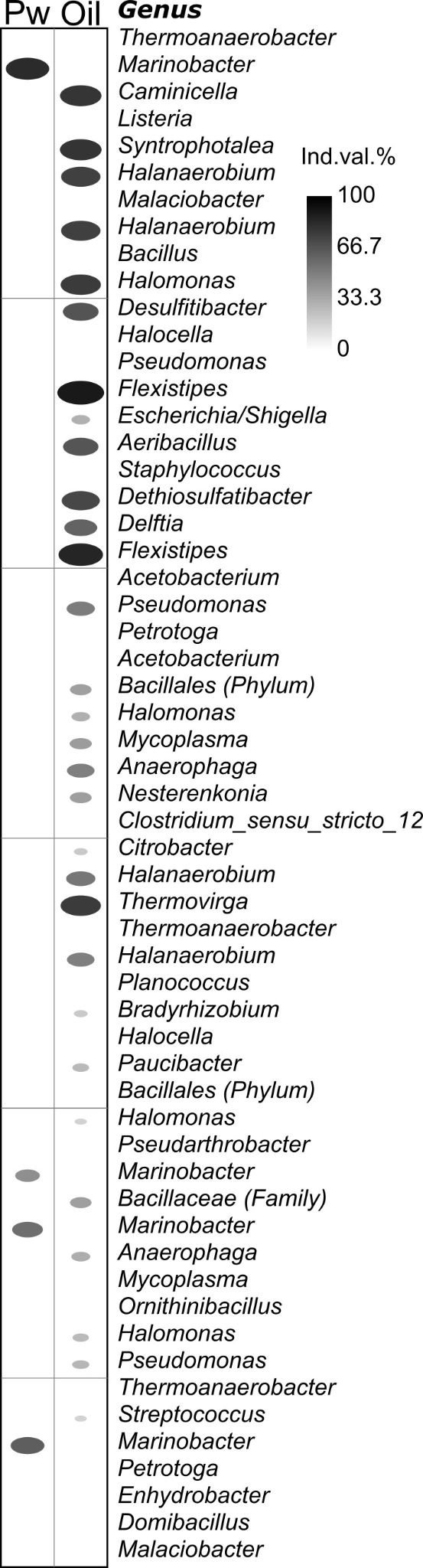

**Figure S1: Indicator species analysis IndVal of the most abundant taxa (cutoff 10 reads).** The taxa that show an association (a circle) have a p-value below 0.05 means the observed association is unlikely to have occurred by chance and is statistically significant. The score is expressed in percentage where 100% means the taxon is both highly specific to and frequent in the sample, making it a strong indicator for that group.

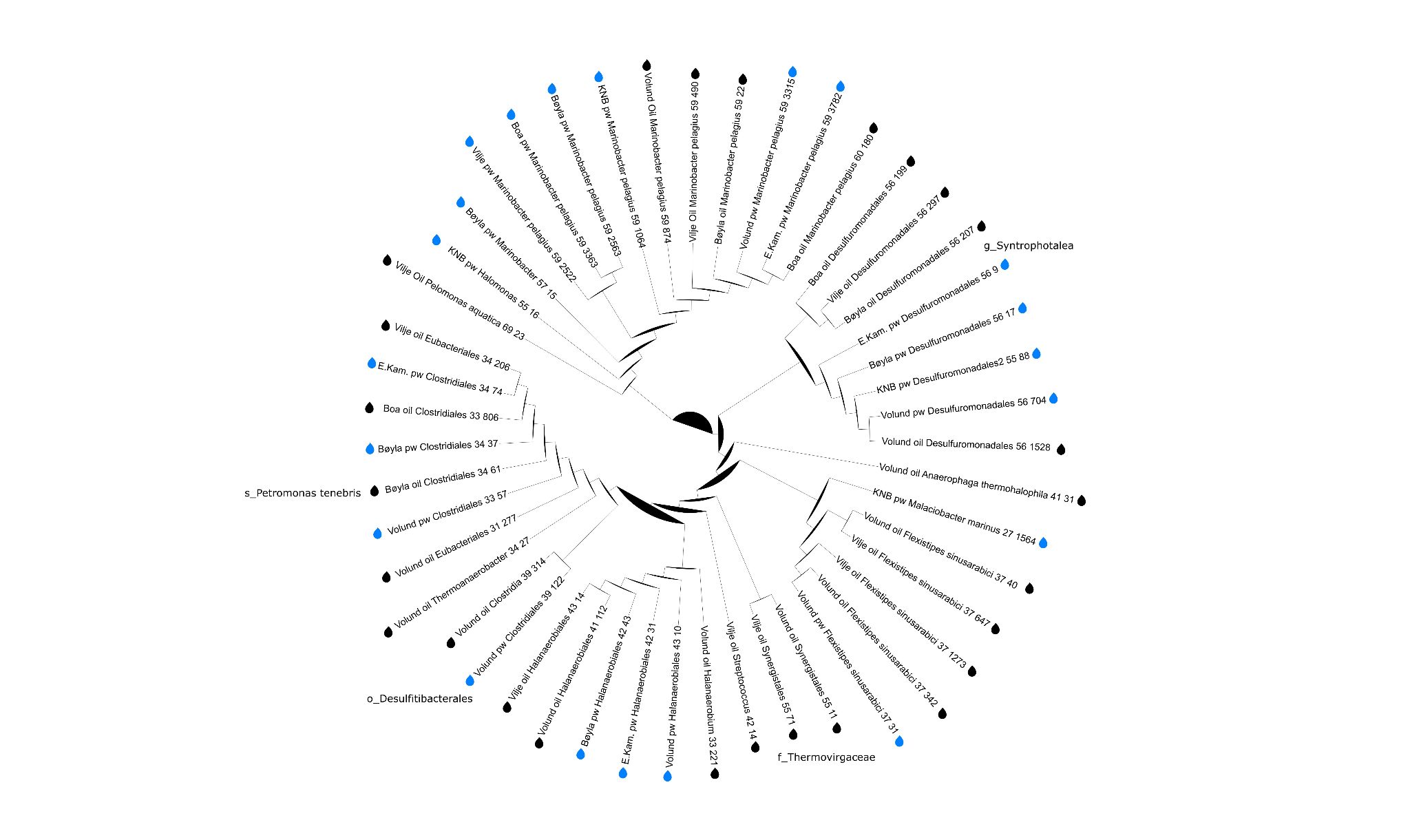
**Figure S2: Phylogenetic tree of metagenome-assembled genomes (MAGs).** The drop shows whether the MAG is originating from production water (pw) or oil samples. The node ID was added in the case it provided more information on the taxonomic assignment.

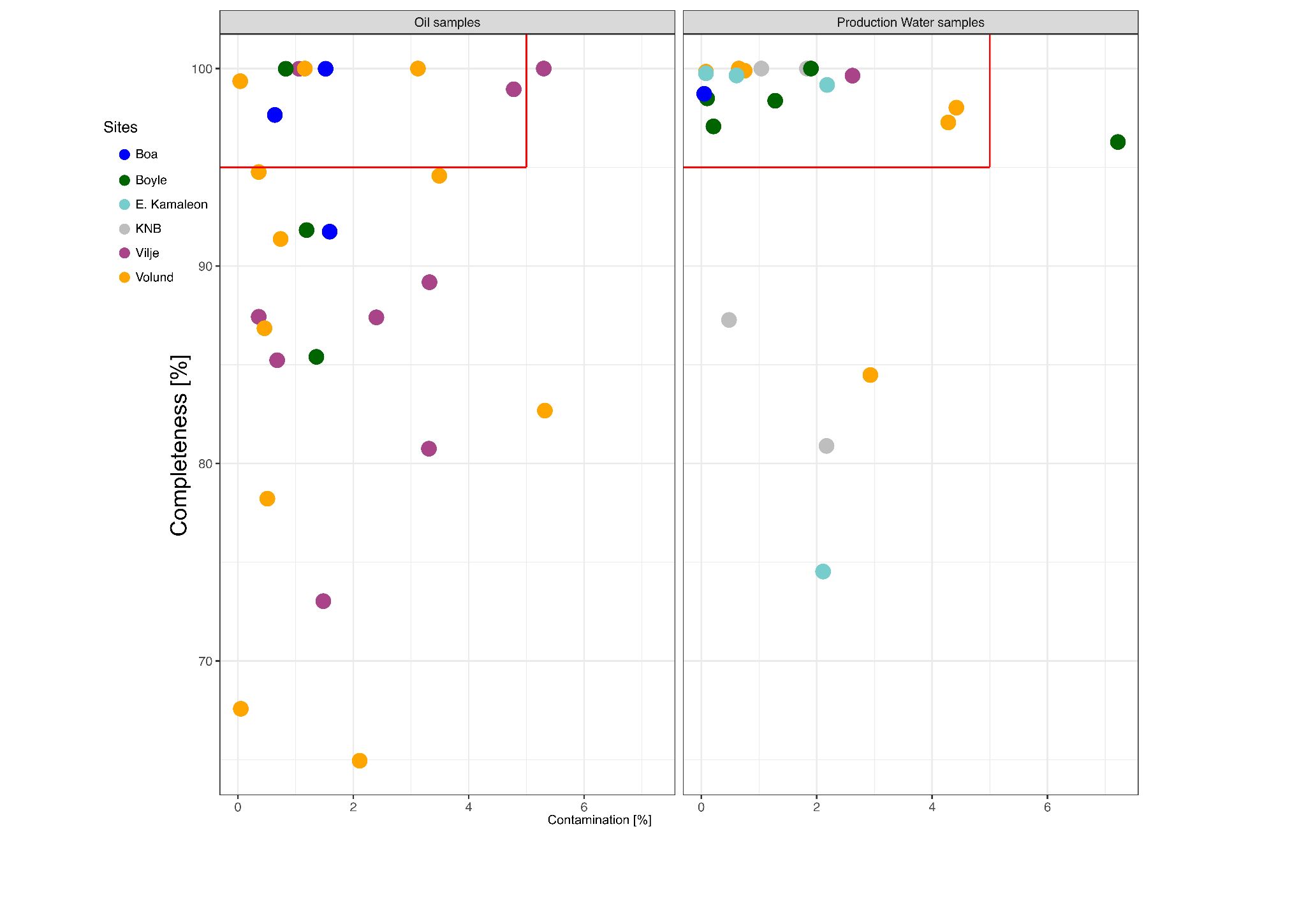

**Figure S3:**  **Completeness and contamination of MAGs from oil and production water samples.** The red rectangle highlights MAGs with a minimum of 95% completeness and a maximum of 5% contamination.

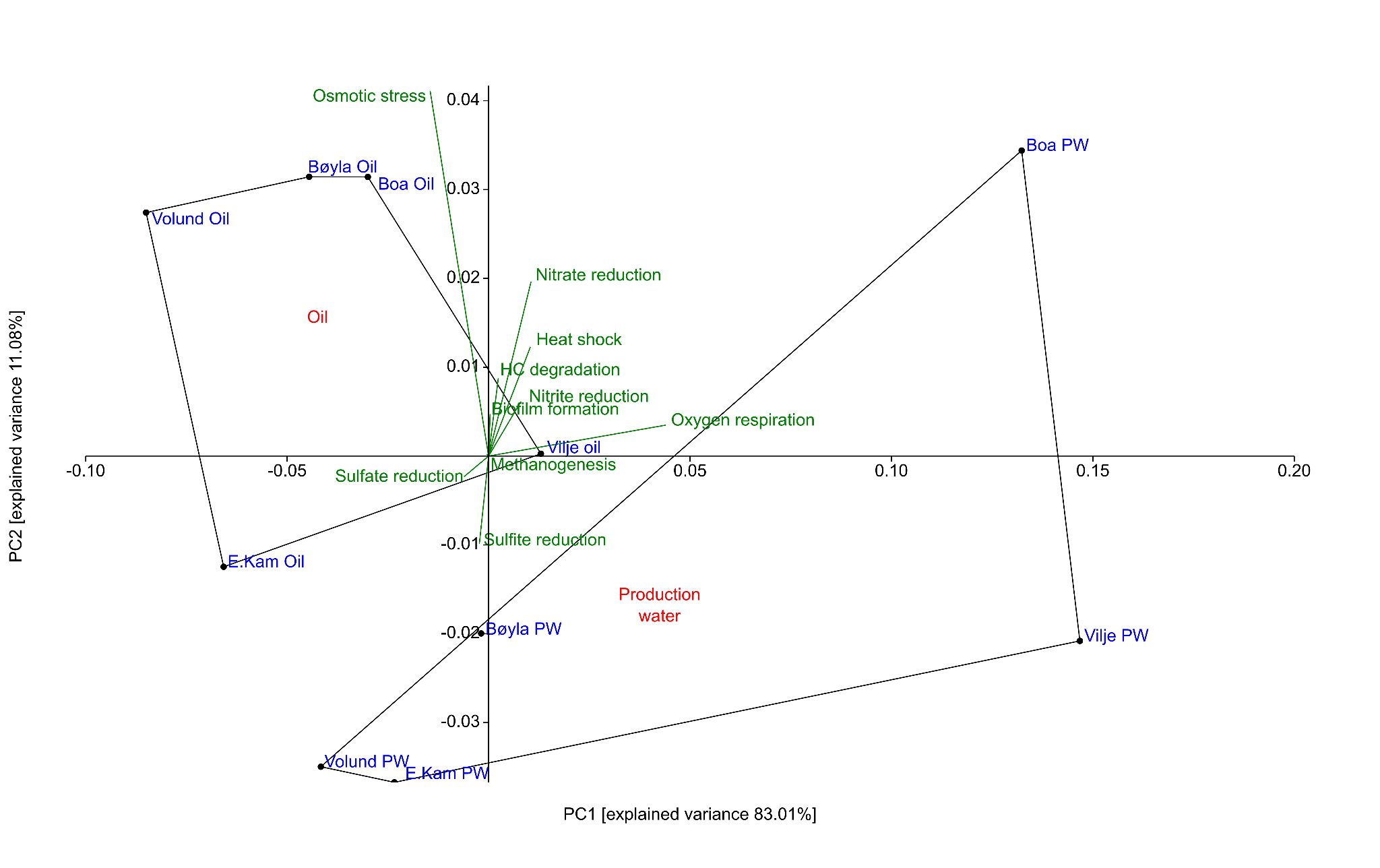

#### **Figure S4: PCA biplot of the ORFs of interest obtained from the samples expressed in percentage of gene hits of the total number of ORFs.** The green arrows indicate the direction towards where the metabolic functions are found.

**Supplementary tables:**

All tables are provided as a separate file (supplementary tables.xlxs).

**Table S1:** Permanova results, in grey the samples that showed no difference

**Table S2:** Selected open reading frames (ORFs) associated with proteins of interest from the BlastP data table.

**Table S3:** Number of ORFs matching the associated pathways and proteins of interest.

**Table S4:** Metadata of the assembly of metagenomic reads into scaffolds for the eleven samples whose libraries were successfully sequenced.

**Table S5:** Coverage and breadth of extended *rpS3 gene* sequences in eleven metagenomic samples and GTDB taxonomy (Parks et al. 2022) of *rp3 gene* sequences.

**Table S6:** Metadata on the *de novo* assembly of the 47 metagenome-assembled genomes obtained from the eleven metagenomic assemblies.

**Table S7:** InStrain (Olm et al. 2021) comparison between the strain profiles of 47 MAGs in the eleven metagenomes.
